# Microdiversity drives resilience and niche partitioning in the bloom forming cyanobacteria *Limnospira platensis*

**DOI:** 10.64898/2026.10.22.760931

**Authors:** A. Le Vern, C. Duval, J. Frère, J.P. Bouly, M. Troussellier, M. Hugoni, H. Agogué, D. Jézéquel, B Marie, C. Bernard, S. Halary

## Abstract

The intraspecific evolutionary mechanisms of adaptation in blooming cyanobacteria, particularly those occurring on small spatial and temporal scales, remain poorly understood. Here, we used the unique properties of Lake Dziani Dzaha, which hosts a persistent population of *Limnospira platensis* inhabiting distinct macro-habitats, to explore the spatial and temporal dynamics of microevolution. The Lake Dziani Dzaha (Mayotte), very stable at least since 2010, has experienced a seismo-volcanic crisis from May 2018 to November 2019, characterized by CO₂ outgassing that led to the mixing of the initially seasonally stratified water column, as well as changes in numerous physicochemical parameters. Phylogenomic analysis of 108 strains isolated during sampling campaigns in 2010, 2014, 2022, and 2024 revealed a population that remained highly phylogenetically cohesive over time. In 2022, genomic expansion was observed, characterized by an increase in genome size due to the proliferation of insertion sequences and a subsequent loss of adaptive genes. By 2024, a rapid recovery of the 2010-2014 genetic content took place through multiple independent events of horizontal gene transfer. At the spatial scale, although no significant differences in gene content were detected, distinct subpopulations were identified based on allele frequencies. Some alleles were specifically fixed and subject to different selection pressures among the subpopulations, suggesting the existence of distinct ecotypes. Overall, this study highlights the role of microdiversity in rapid resilience of blooming cyanobacteria populations within dynamic environments and in their adaptation through niche partitioning.

## Introduction

In bacteria, microdiversity is considered a fundamental feature of adaptation. Defined as the coexistence of genomically and physiologically distinct individuals of the same lineage, microdiversity enables populations to cope with spatial and temporal environmental variations (Moore *et al*., 1995, Fuhrman & Campbell, 1998). This phenomenon has been thoroughly studied in marine picocyanobacteria *Prochlorococcus* and *Synechococcus*, two dominant primary producers that have colonized most photic zones across the world’s oceans, for which a wide diversity of genotypes has been clustered into functionally and ecologically coherent groups, referred to as ecotypes (Biller *et al*., 2015, Rocap *et al*., 2003).The functional traits that differentiate these ecotypes have been linked to a wide range of environmental parameters, including temperature, nutrient and trace metal availability, as well as light intensity and quality (Moore *et al*., 1995, Martiny *et al*., 2009, Grébert *et al*., 2018). Consequently, ecotypes preferentially occupy distinct ecological niches, with their relative abundance determined by gradients along lines of latitude, depth, distance from continental margins, or across seasons (Flombaum *et al*., 2013, Mattei *et al*., 2025). This structure reflects a process of niche partitioning, in which ecotypes have diverged to exploit complementary portions of the environmental landscape, thereby reducing direct competition (Johnson *et al*., 2006).

Bloom-forming cyanobacteria pose a serious threat to lake ecosystems and public health, due to their ability to proliferate near the surface (Ibelings *et al*., 1991) and for some genotypes to produce harmful secondary metabolites (Wu *et al*., 2014, Halary *et al*., 2023). Although cyanobacterial blooms are becoming increasingly frequent, driven by watershed anthropization and global change (O’Neil *et al*., 2012), their population dynamics remain poorly understood. Microdiversity may play a major role in bloom development (Wu *et al*., 2024). Thus, subpopulations with distinct OTUs succeed in one another over time and space during a bloom (Briand *et al*., 2009, Banjeri *et al*., 2019). Likewise, several genotypes may coexist within a single bloom, exhibiting marked differences in both global gene content and their capacity to synthesize secondary metabolites (Halary 2023, Yéprémian *et al*., 2017). Although ecotypes of Microcystis have been characterized based on phosphate tolerance and thermal adaptation (Jackrel et al., 2019; Kuipers et al., 2025), these represent the few documented examples to date, as identifying ecotypes is challenging when the adaptive traits or niche differences underlying population differentiation are unknown.

Indeed, bloom-forming cyanobacteria generally thrive on the surface of small shallow lakes, where the physico-chemical parameters are virtually isotropic, which represents a major difference from the ocean. It is therefore thought that the high level of genotypic diversity observed there reflects a significant heterogeneity of micro-niches (Halary *et al*., 2023, Huré *et al*., 2026), which can sometimes be reduced to the mere proximity of a particle of organic matter (Bobay, 2020), and which is difficult to characterize using conventional approaches in microbial ecology. Another challenge in microdiversity studies lies in defining biologically meaningful taxonomic units based on genomic variation. For example, Prochlorococcus constitutes a diverse clade within which distinct ecotypes may exhibit average nucleotide identity (ANI) values well below the conventional 95% threshold used for species delineation (Zhaxybayeva et al., 2009; Richter & Rosselló-Móra, 2009). Similarly, the taxon Microcystis aeruginosa has been associated with numerous distinct genospecies within the genus, many of which can coexist within the same ecosystem (Huré et al., 2026). Under these circumstances, the contribution of microdiversity to ecological processes may be blurred by variation that reflects interspecific rather than intraspecific diversity (Bobay, 2020).

To investigate the ecological significance of microdiversity in blooming cyanobacteria while reducing these confounding effects, we focused on the *Limnospira* (Oscillatoriales) population inhabiting the atypical environment of Lake Dziani Dzaha (Mayotte, France). This small thalassohaline volcanic crater lake, covering an area of 25 ha, is located in the Comoros archipelago (Zinke et al., 2003; Cadeau et al., 2020). Its average depth is 4 m, although a circular pit in its eastern part reaches 18 m. The *Limnospira* from the lake belong to clade I of the species *L. platensis* (Roussel *et al*., 2023); they formed a persistent bloom on the water column surface, dominating an extremely high phytoplanktonic biomass (up to 800 µg Chl*a* L^− 1^ in April 2014), but have also colonized the biofilms that develop on hard substrates and the green biotic layers of the stromatolites along the shores, although they do not contribute to their formation (Bernard *et al*., 2019, Gerard *et al*., 2018). Since survey began in 2010, this lake used to altern between stratified and unstratified states depending on the season, with a chemocline at about 2 m. Although surface water was always highly oxic, a steep oxygen gradient developed with depth, reaching anoxia at the chemocline. From May 2018 to November 2019, a seismic crisis triggered a magmatic CO₂ discharge from the pit, that disrupted the lake’s dynamics, characterized notably by the loss of seasonal stratification, a drop in pH from 9.5 to 8.1 (Feuillet *et al*. 2021, Sarazin *et al*. 2021, Cadeau *et al*. 2022).

All these characteristics make Lake Dziani Dzaha a natural laboratory of interest for reverse ecology, enabling the study of the spatial and temporal dynamics of microdiversity within a coherent cyanobacterial population. This study aims to test the hypotheses of the impact of microdiversity on resilience and niche partitioning of blooming cyanobacteria.

## Materials and Methods

### Sampling and *Limnospira* strains isolation

The isolation of *Limnospira* strains is based on four sampling campaigns conducted in September 2010 (DZ10-09), April 2014 (DZ14-09), June 2022 (DZ22-06) and June 2024 (DZ24-06) (see Table. S1 for details). Water samples were collected at three stations (ET, St3 and CLB) (Fig. 1A) at various depths (0.25, 1.0, 2.5, 4.0, 16.0 and 17.25 m) using a 2.5 L horizontal Niskin bottle and were processed in the field laboratory within two hours. Biofilms were collected from the surface of buoy B1 (DZ24-06) and stromatolite S4 (DZ14-04) (Fig. S1A-D), and samples of surrounding water were collected for the stromatolites and biofilms from the DZ24-06 campaign. Strains from the stromatolite biotic layers of the DZ24-06 campaign were isolated from fragments of the S2 and S3 stromatolites. Prior to isolating the S2 and S3 stromatolites, the fragments were carefully brushed with culture medium to minimize potential contamination by other microorganisms present on their surfaces. The inner face of each fragment was then placed onto the culture medium, following the protocol described below. Finally, strains were also isolated from surface water surrounding S2, S3 and B1 (DZ24-06).

**Figure 1.**
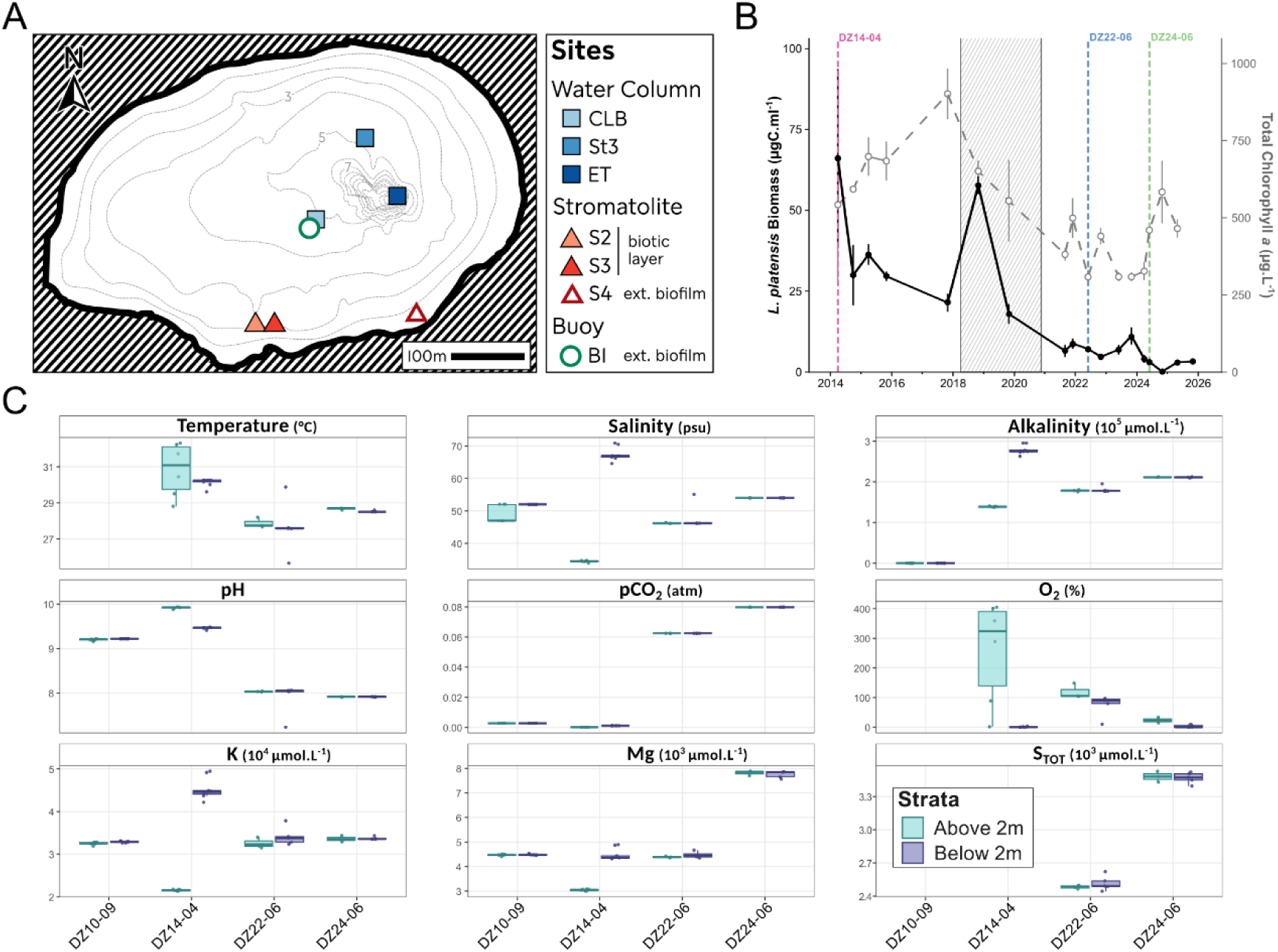
Overview of Lake Dziani Dzaha sampling sites, and temporal characteristics of the water column. **A** – Bathymetric map of the lake and locations of the sampling stations (adapted from Cadeau et al. 2022) **B** – Temporal monitoring of Limnospira platensis biomass (black bold line, primary y axis) and total chlorophyll a concentration (grey dashed line, secondary y axis) in the surface layer of the water column (above 2m). The colored dashed vertical lines indicate the sampling campaigns (DZ10-09, DZ14-04, DZ22-06, DZ24-06), and the hashed grey zone represents the seismic crisis event. **C** - Temporal variation in physicochemical parameters above and below the chemocline, including temperature, salinity, alkalinity, pH, partial pressure of CO₂ (pCO₂), and concentrations of dissolved oxygen (O₂), potassium (K), magnesium (Mg) and total sulfur (S_TOT_).

All samples were inoculated on semi-solid (5 g L^−1^ of agar) *Spirulina* medium (Aiba & Ogawa, 1977; Hamlaoui *et al*. 2022). The Petri dishes were half-covered with aluminum foil and incubated from 24h to 48h at 25°C, to induce phototactism. *L. platensis* isolates were obtained by repeatedly transferring single filaments from a solid medium to a liquid medium under an inverted microscope (Nikon ECLIPSE TS100), using a tapered Pasteur pipette. Growing clones were transferred to 25 cm² Falcon® culture flasks containing 10 mL of *Spirulina* medium and maintained in the Paris Museum Collection (PMC, MNHN), at 25 °C under LED illumination (10 μmol photons.m⁻².s⁻¹) with a 13:11 h light photoperiod.

Vertical profiles of physical and chemical parameters along the water column were performed as described in Cadeau *et al*. 2022 and *Limnospira platensis* biomass was calculated as described in Menden-Deuer & Lessard 2000.

### Genome acquisition and comparative genomics

Cultures were maintained for 18 days and stopped at an OD_750nm_ of 0.4. DNA extraction was performed using the Qiagen DNeasy® Powerlyzer® Powersoil® bead beating kit (QIAGEN) and sequenced using a shotgun metagenomic approach (paired-end 2×150 bp) on the Illumina NovaSeq platform (Genoscreen®, Lille). Raw reads were quality-trimmed with Trimmomatic v0.39 (Bolger *et al*., 2014) using default parameters. Assemblies were performed using SPAdes v3.15 (Bankevich *et al*., 2012) and *Limnospira* Metagenome-Associated Genomes (MAGs) were extracted as described in Halary *et al*. (2022). After quality assessment using CheckM2 (Chklovski *et al*., 2023), MAGs displaying a completeness < 98% and a contamination > 3% were discarded. Two closed relative *Limnospira platensis* genomes were added to the analysis as references, namely SAG85.79 and TJSD092. Pairwise Average Nucleotide identity (ANI) comparisons were calculated using fastANI (Jain *et al.,* 2018). For each genomes/MAGs, open reading frames prediction was performed using Prokka v1.14.6 (Seemann 2014). Differences in genome sizes and number of coding sequences were tested first by ANOVA and by Tukey *post hoc* test when applicable. The number of pseudogenes was estimated using Pseudofinder (with NR as the reference database) (Syberg-Olsen *et al*., 2022). Only pseudogenes supported by an identifiable disrupted ORF were retained. Insertion Sequences (IS) were also surveyed using ISESCAN (Xie & Tang, 2017). Coding sequences were functionally annotated using EggNOG-mapper v2.1.12 (Huerta-Cepas *et al.,* 2019). Orthologous groups were defined using OrthoFinder v.3.1.2 (Emms & Kelly, 2019) and used as input to create orthologs presence/absence matrix. For each orthogroup, protein sequences sharing 100% identity over their full length were clustered to determine the presence of distinct protein variants.

### Phylogenomics analysis

A concatenation of all alignments from genes belonging to the core genome was produced by PANAROO (using MAFFT aligner and –clean-mode option ‘strict’). Recombined regions were filtered using gubbins (Croucher *et al*., 2015). The resulting alignment served as input for a phylogenetic inference performed by IQTREE2 (GTR+F+R4 model, with 1000 ultrafast bootstraps and bootstrap correction -bnni option) (Minh *et al*., 2020).

### Multivariate Statistical Analyses

All statistical analyses were performed in R (v2026.1.1.403; R Core Team, 2026). Principal Coordinate Analyses (PCoAs) were computed from Jaccard distance matrices using vegan (v2.7-2; Oksanen *et al*., 2025) and stats (R Core Team, 2026) packages. Groups were defined using the NbClust package (v3.0.1; Charrad et al., 2014), with a maximum of five clusters and the “alllong” index. Population gene content structure was assessed by stratified permutational multivariate analysis of variance (PERMANOVA). Environmental variables were tested both jointly and individually. Homogeneity of multivariate dispersions was assessed using permutest, followed by ANOVA to compare dispersion among habitat groups. Pairwise comparisons were adjusted using the Benjamini–Hochberg (BH) correction.

To investigate the effects of biotic and abiotic parameters on population genomic variation, pairwise Pearson correlations were first used to identify highly colinear variables, with variables showing strong correlations (> 0.7) being represented by a single proxy variable in subsequent analyses. Distance-based redundancy analysis (db-RDA) was performed using the capscale function to assess relationships between environmental variables and population gene content. Mantel tests (999 permutations) were used to evaluate associations between the Jaccard distance matrix and categorical distance matrices (sampling site, sampling campaign, genomic cluster).

Indicator genes were identified using the indicspecies package (De Cáceres *et al*., 2009). Only orthologs displaying a specificity (A; the frequency of the gene within a group) ≥ 0.7 and a fidelity (B; the proportion of genomes in that group carrying the gene) ≥ 0.5 with a padj (BH) < 0.05 were kept as indicators.

### Homologous recombination rates across *Limnospira* genome

All genomes were aligned to the genome of the most central branch of the phylogenomic tree, and a multiple alignment of core genes was obtained using snippy (Seemann, 2015). A Maximum Likelihood inference was then performed using IQTree2 (Minh *et al*., 2020) with the following parameters: GTRG, -B 1000 -bbni. The resulting tree was rooted on the reference. The reference was removed from both the tree and the alignment before they were used as input for ClonalFrameML (Didelot *et al*., 2007) with branch-specific parameter estimation (-- embranch, 100 EM simulations). The required transition-transversion ratio was estimated from IQTree2 results. Differences of resulted indices among different groups (Year, PcoA Clusters, Habitat) were investigated using Kruskall-Wallis and *post-hoc* Dunn tests.

### Differentiation and selection analyses

Single copy genes of the core genome were used to calculate fixation and selection indices. For each, a multiple alignment of protein-coded sequences was performed by MAFFT (Katoh *et al*. 2013) in *localpair* mode. Corresponding nucleic sequences were then aligned based on the corresponding multiple amino-acid sequence alignment using pal2nal (Sumaya *et al*., 2006). F_ST_ among PCOA clusters were calculated using fst.hudson function from R-PopGen package (Lesturgie *et al*., 2022), with 999 resampling, and p-values adjusted using BH correction. Genes displaying a F_ST_ > 0.25 with p-adj < 0.05 were considered as outliers. Selection analyses were performed using HyPhy (v2.5.99). Branch-level selection was tested with BUSTED-PH (with synonymous rate variation) to detect gene-wide and foreground-specific positive selection. Site-level selection was assessed using FEL (pervasive positive or purifying selection at codons) and MEME (episodic diversifying selection). After p-values were corrected with BH, genes were classified as foreground-specific positively selected, site-selected, under purifying selection, or showing no detectable selection signal (see Table. S2).

## Results

### Population decline associated with environmental change

The biomass of *L. platensis* fluctuated between 2014 and 2018, with values ranging from 21. 5 ± 2.8 µg C L^−1^ to 66.3 ± 27.7 µg C L^−1^, and steadily decreased since 2018, reaching values close to 1.5 µg C L^−1^ by the end of 2024. The total chlorophyll *a* content, which integrates other phototrophic organisms, has also fallen after a peak of 901.7 ± 80.6 µg L^−1^ in 2018 but rebounded significantly in early 2024 (Fig. 1B).

Environmental parameters revealed changes in water composition through depth or time (Fig. 1C). The stratified periods could be identified by the variations between the surface and the hypolimnion of temperature (30.84 ± 1.48 °C vs 30.14 ± 0.21 °C), salinity (34.45 ± 0.30 psu *vs* 67.39 ± 2.02 psu) and dissolved oxygen (257.40 ± 171.82 % *vs* 0.67 ± 0.92 % respectively) during DZ14-04 campaign, although these factors quickly fluctuated within the first 2 m from the surface.

The differences between parameters above and below the chemocline during the DZ14- 04 cruise (e.g., temperature, alkalinity, pH)—which are characteristic of stratification during this season—were no longer apparent in subsequent campaigns (DZ22-06 et DZ24-06). Moreover, several environmental parameters changed significantly and became homogeneous across the surface layer between 2014 and 2024: Surface-water pCO₂ increased 570-fold (from 1.4·10^−5^ atm to 8.0·10^−3^ atm), while dissolved oxygen concentrations decreased 11-fold ± 8.63 (from 257.40 ± 171.82 % to 23.27 ± 9.39 %) and pH declined from 9.92 ± 1.97·10^−2^ to 7.91 ± 4.62·10^−3^. Concentrations of K^+^ (21,550 ± 184 µmol·L^−1^ to 33,593 ± 750 µmol·L^−1^) and Mg^2+^ (3,045 ± 36 µmol·L^−1^ to 7,811 ± 102 µmol·L^−1^) likewise both increased. Finally, total sulfur concentration increased from 2.5 to 3.8 µmol·L⁻¹ between 2022 and 2024, the only two years for which measurements were available (Fig. 1C).

### High genomic similarity despite gene-content variations

Of the 121 strains isolated and sequenced, 108 genomes were kept for analysis after the quality control (Table S1). All genomes shared the same GC content (45.0%), and showed limited variation in genome size (5.82 ± 0.13 Mbp), protein-coding gene number (4,831 ± 58.06) and ortholog count (4,781 ± 55.86; Table S1). Despite this overall genomic similarity, the pangenome comprised 6,642 orthologous groups, of which 3,837 (57.8%) belonged to the core genome and 2,805 (42.2%) to the flexible genome. Notably, 999 orthologous groups (15.0% of the pangenome) were singletons, *i.e,* detected in only one genome (Fig. S2). Pairwise average nucleotide identity (ANI) further confirmed the high genomic similarity among L. platensis isolates from Lake Dziani Dzaha, with a mean of 99.69 ± 0.09% after excluding self-comparisons. ANI values were slightly lower in comparisons with the outgroup strains SAG85.79 and TJSD092 (99.53 ± 0.01%; Fig. S3). A PERMANOVA analysis identified sampling campaign as the only tested factor significantly associated with ANI-based genomic distances (R² = 0.34, p < 0.001).

### Temporal and spatial mixing dominates the phylogenomic structure

The phylogenomic tree obtained from the core genome expurged from most of the recombination signal is shown in Fig. 2A. Strains from different campaigns and different habitats were distributed and mixed all along the tree, at the exception of 3 strains isolated in 2014 from the biofilm S4, and those isolated in 2010, from the water column. These oldest clades were both distant from the root. Both 2022 and 2024 strains can be clustered into year-specific clade, which can be as large as 20 leaves. Many clades supported by a robust bootstrap > 80 gathered strains sampled from different times. Phylogenomic inference based on raw core gene multiple alignments displayed a near identical topology (data not shown).

**Figure 2.**
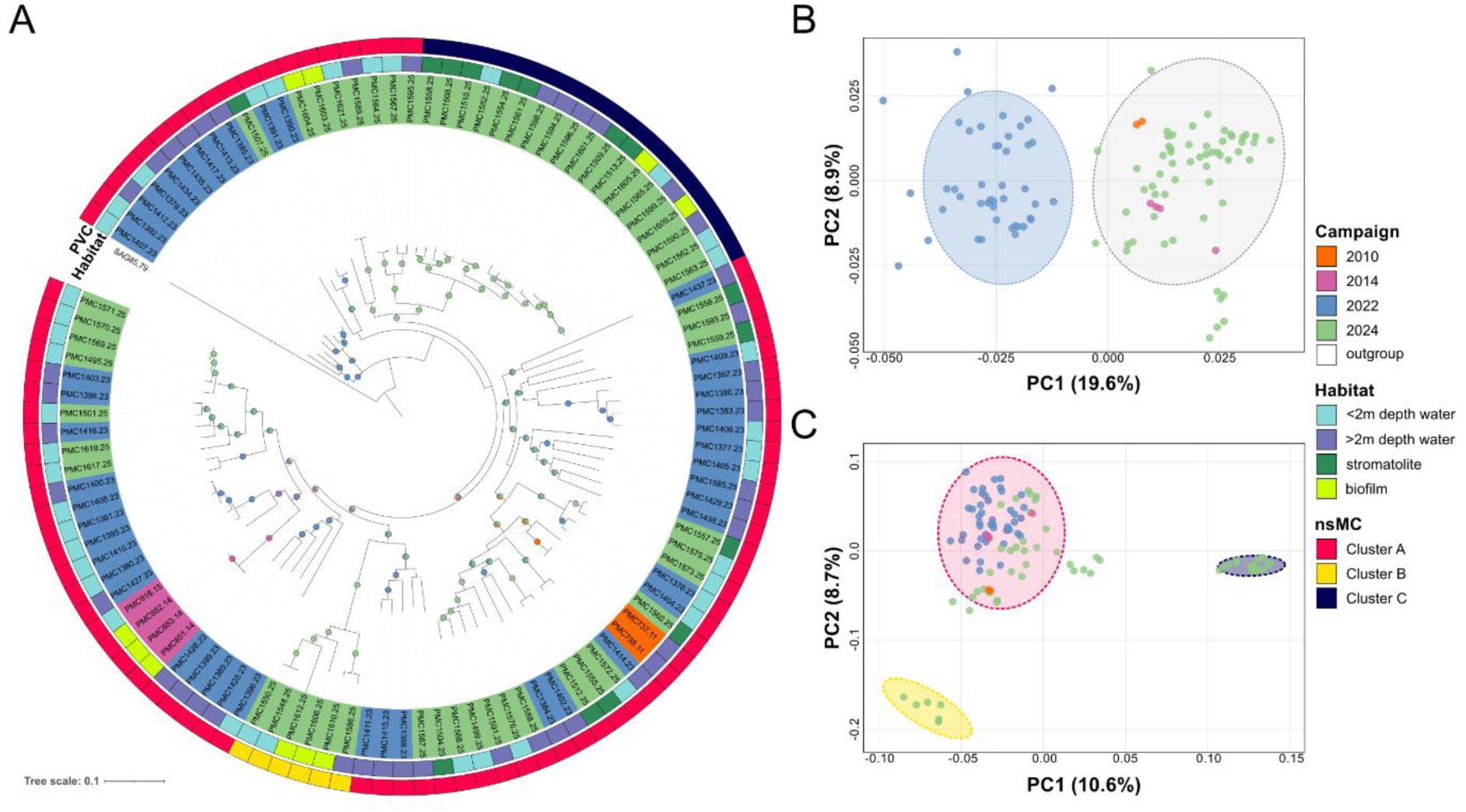
Phylogenomic tree of Limnospira platensis from Lake Dziani Dzaha and genomic structure of the population. **A** - Maximum-likelihood phylogeny inferred from the recombination-filtered core genome. Leaves are colored following the strain sampling year. Bootstrap values >80% are represented by circles at the corresponding branches, with colors indicating the sampling campaign of the strains belonging to the supported clade. The inner circle (Habitat) provides information on the ecological origin of the strains (surface water <2 m, deep water >2 m, stromatolite, or biofilm). The outer circle indicates the protein variants cluster leaves belong (from C.). The tree was rooted on the L. platensis (Clade I) strain SAG85.79 isolated from Natron lake (Chad). **B** - Principal Coordinate Analysis (PCoA) based on Jaccard distances among gene contents (presence/absence). Each point represents a strain, colored according to the sampling year. C- PCoA based on the protein variant composition of single-copy genes in the core genome. For both PCoAs, ellipses represent 95% confidence intervals for statistically distinct groups.

### *Limnospira* population structure at the gene content and protein variant levels

The Principal Coordinates Analysis based on whole gene contents discriminated genomes into two clusters (11/27 NbClust validated indices and PERMANOVA, R²=12.30, p<0.001), with PC1 and PC2 accounting for 19.6% and 8.9% of the total variance, respectively (Fig. 2B). The first cluster gathered all strains from 2022, while the second grouped those from 2010, 2014, and 2024. The ‘2022’ cluster displayed a greater dispersion compared to the other one (0.345 *vs.* 0.308; betadisper permutation test F=5.40, p=0.016). The greater gene content heterogeneity of the strains isolated in 2022 was also supported by a greater size of its pangenome (Fig. S2A). The analysis of physicochemical gradients, from which pCO_2_ was discarded due to limited number of measurements, revealed strong collinearities among variables, forming two groups: one associated with salinity, alkalinity, temperature, Mg and K, and the other with pH, dissolved O₂ and *L. platensis* biomass (Fig. S4A). Salinity and pH, chosen as representative variables (Fig. S4A), accounted for 16.64% and 5.54% of the total variance, respectively (p < 0.001, Fig. S4B). Both significantly participated in the Distance-based redundancy analysis (dbRDA) to the segregation of strains from 2022 to all others on the axis 1, similarly to the PCoA based on gene contents (Permutation tests, pval < 0.05). But strains from 2010 and 2014 were also separated on the axis 2 from the 2024 ones.

PcoA based on the presence/absence pattern of nonsynonymous mutation in core genes (nsM)identified three clusters (23/27 validated NbClust indices, PERMANOVA R² = 16.83%, p < 0.001), named nsM Cluster (nsMC) A, B and C (Fig. 2C). Group nsMC A comprised most of the strains (76.8%) from all habitats and sampling campaigns (Fig. 2A). nsMC B (5.6%) gathered strains from water column and biofilms, while nsMC C (17.6 %) grouped strains from water column, biofilms and stromatolites, both consisted exclusively of strains from 2024. Each nsMC displayed a distinct dispersion (pairwise PERMANOVA, padj<0.003), with the highest observed in nsMC A (0.41), followed by nsMC B (0.29) and nsM C (0.11).

### Temporal divergence in genome size, gene content and pseudogenisation

Indicator gene analysis identified 69 orthologs as markers for the ‘2022’ cluster, and 168 for the ‘2010’s/2024’ cluster (Figs. 2B, 3A). For the first, only six genes over 69 obtained a functional annotation (Table. S3), including three transposases and a poorly characterized protein. The last two genes coded for proteins of the chaperone-usher (CUP) system, which is essential for pilus assembly involved in cell-to-cell adhesion (Schuergers & Wilde, 2015). In contrast, 45 indicator genes over 168 of the 2010’s/2024 cluster were functionally annotated (Table. S3), including 21 coding for proteins involved in information storage and processes, cellular processes, or metabolism. Besides functional gene content, significant differences were also found at the genome size and protein-coding genes number levels among sampling campaigns, with strains from ‘2022’ cluster displaying the largest genomes (µ_2022_=5.94 ± 0.05 *vs.* µ_2010s_= 5.72 ± 0.09 and µ_2024_= 5.75+-0.11) and the lowest number of coding genes (µ_2022_ = 4797.45 ± 57.72 *vs.* µ_2010s_ = 4860 ± 77.41 and µ_2024_ = 4853.45 ± 42.65, Fig. 3B). Genome size was strongly associated with insertion sequence (IS) abundance (R² = 0.771, *p* < 2.2 × 10⁻¹⁶), with the data revealing two distinct groups of genomes (Fig. 3B). Genomes smaller than 5.82 Mbp harboured a mean of 73.12 ± 6.35 IS, whereas larger genomes contained 124.05 ± 13.58 IS.Moreover, intact genes (*i.e.,* not pseudogenized) were more abundant in 2010s (median=4390) and 2024 (median=4400) genomes than in 2022 ones (median=4310, padj < 0.05), whereas truncated pseudogenes which are the most frequent category found displayed the inverse pattern (median_2010s_=395, median_2022_=445, median_2024_=400, Fig. 3B).

**Figure 3.**
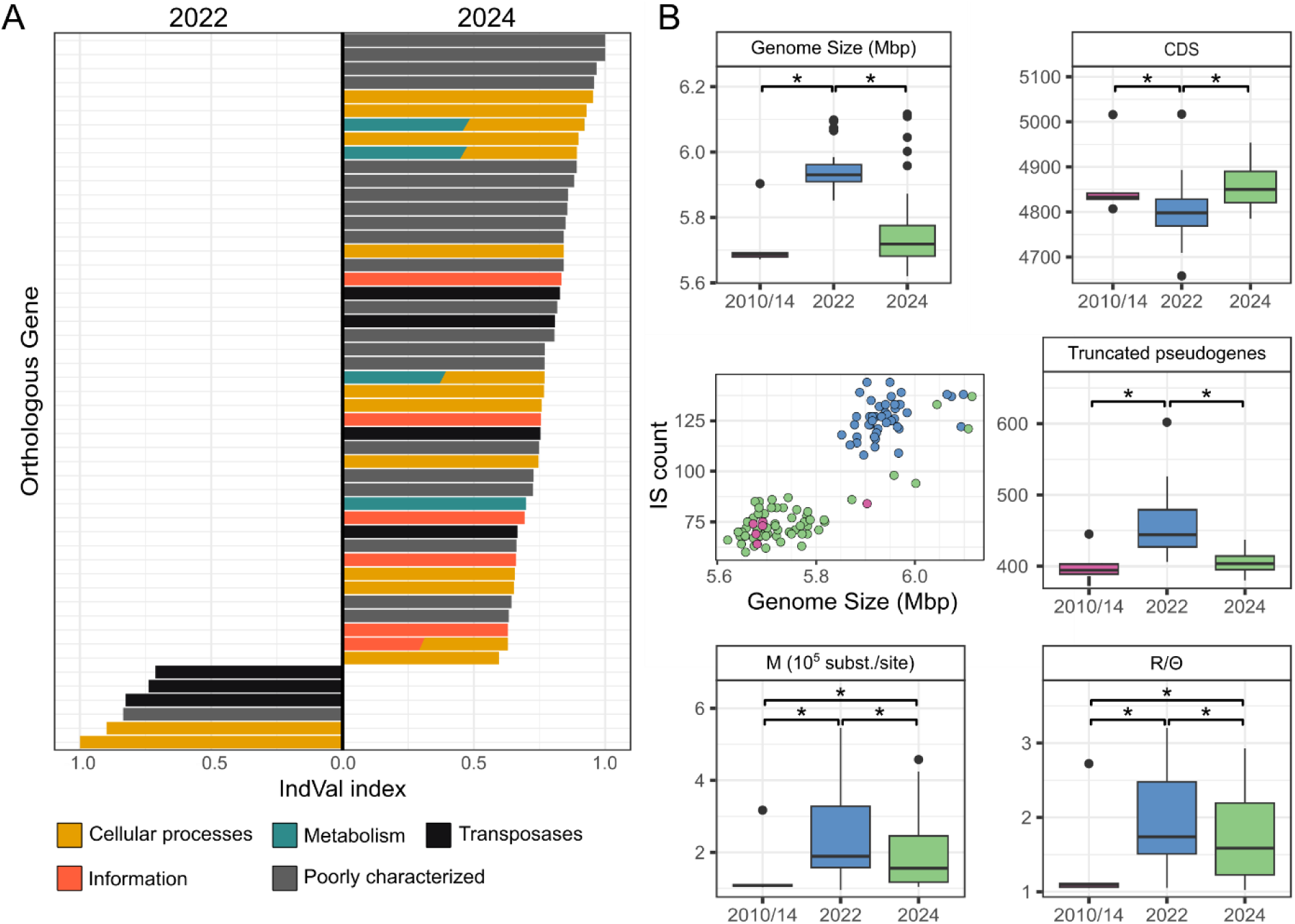
Temporal dynamics of genomic differentiation and genome architecture. **A** - Indicator genes identified by IndVal analysis across 2022 and 2024. Bars indicate IndVal values for orthologous genes displaying a specificity A>0.7 and a fidelity B>0.5 (p < 0.05), colored following the COG functional categories. **B** - Boxplots and scatterplots illustrating temporal changes in genome architecture and population genomic parameters. From left to right and top to bottom: genome size; number of predicted coding sequences (CDSs); relationship between insertion sequence (IS) abundance and genome size; number of short pseudogenes; the per-branch substitution rate (M); and the recombination-mutation ratio (R/θ).

To characterize the evolutionary dynamics underlying the observed population structure, we assessed the temporal signal and quantified the relative contributions of mutation and recombination to genomic diversification. The absence of a significant correlation between genetic distance and sampling date (R² = 0.024, TempEst) indicates that the population lacks detectable molecular clock signal, precluding estimation of the absolute mutation rate and effective population size. Phylogenomic analyses revealed a highly heterogeneous among-site substitution rate distribution (α = 0.020), violating the homogeneous substitution rate assumption underlying ClonalFrameML (Didelot & Wilson 2015), and likely biasing the estimates of ν and δ toward inflated and deflated values, respectively. For these reasons, interpretation was restricted to recombination-mutation ratio R/θ and the per-branch mutation rate M, which are less sensitive to among-site rate heterogeneity bias as they are primarily informed by the overall density of substitution events rather than their spatial distribution along the alignment (Didelot & Wilson 2015). Both parameters were significantly elevated in 2022 strains relative to 2010/2014 and 2024 (Kruskal-Wallis, Dunn post-hoc, p < 0.05; Fig. 3B).

### Progressive allelic differentiation among co-occurring subpopulations

To identify the molecular signatures underlying the divergence among nsMCs, fixation and selection analyses were conducted on two complementary datasets: first, between 2022 and 2024 genomes within cluster nsMC A (Table. S4), to characterize the genomic recovery following the seismic disturbance event; and second, between 2024 genomes of nsmC A, B and C (Table. S5). Although no significant overall differentiation was detected between nsMC A-2022 and nsMC A-2024 genomes, 34 genes displayed significant allelic fixation between these time points, of which 30 showed detectable signatures of selection in at least one group. Only 5 genes (16.7%) obtained a name after functional annotation. The functional landscape was overrepresented by genes involved in cell wall remodeling and membrane biogenesis (*e.g., rfb*A and *rfb*W, COG category M, 16.7%) and coenzyme transport and metabolism (*e.g., men*D, category H, 10%) (Table. S4).

The comparison between A, B and C revealed a substantial genetic diversification, with a greater F_ST_ value between B and C (0.651, pval < 0.05) than between A-B (0.396) or A-C (0.286). In total, 541 genes showing significant fixation between at least two clusters, distributed throughout the composite genome of PMC 1556.24, the most centrally positioned strain in the phylogenomic tree (Fig. 4A). Of these, 198 displayed signatures of purifying or positive selection in at least one cluster (Fig. 4A, Table. S5), of which 56 (28.3%) received a complete functional annotation with a gene name. Four major metabolic pathways emerge from this functional landscape: photosynthesis, respiration, osmoregulation and oxidative stress response (Fig. 4B). For photosynthesis, *cup*A and *cp1*2 are involved in inorganic carbon uptake, *psb*V and *psb*V2 in photosystem II assembly and adaptation, and *crt*H in carotenoid biosynthesis. All these genes are under selection in nsmC C, with *cp1*2 also showing pervasive positive selection in cluster A. For respiration, *hoxE* and *hoxU* code both for distinct subunits of the bidirectional hydratase complex, whereas *hypA* code for a protein involved in the hydratase maturation. All these genes are under purifying selection in nsMC-A. Two genes known for their implication in osmoregulation (*ggpS* and *proC*) showed selection signatures (positive and purifying, respectively) in nsMC C (*ggp*S, *proc*C). In nsMC-A, *nha*P gene, coding for a protein also involved in osmoregulation were under positive selection. Three oxidative stress response genes were also found to be under selection, namely *hsl*O, involved in protecting thermal unfolding and oxidatively damaged protein from aggregation, in nsMC A, *gor*, involved in glutathione reductase biosynthesis, in nsMC B, and *suf*D, involved in Iron-Sulfur cluster used in PSI, in nsMC C) (see Table S6 for more details and references).

**Figure 4.**
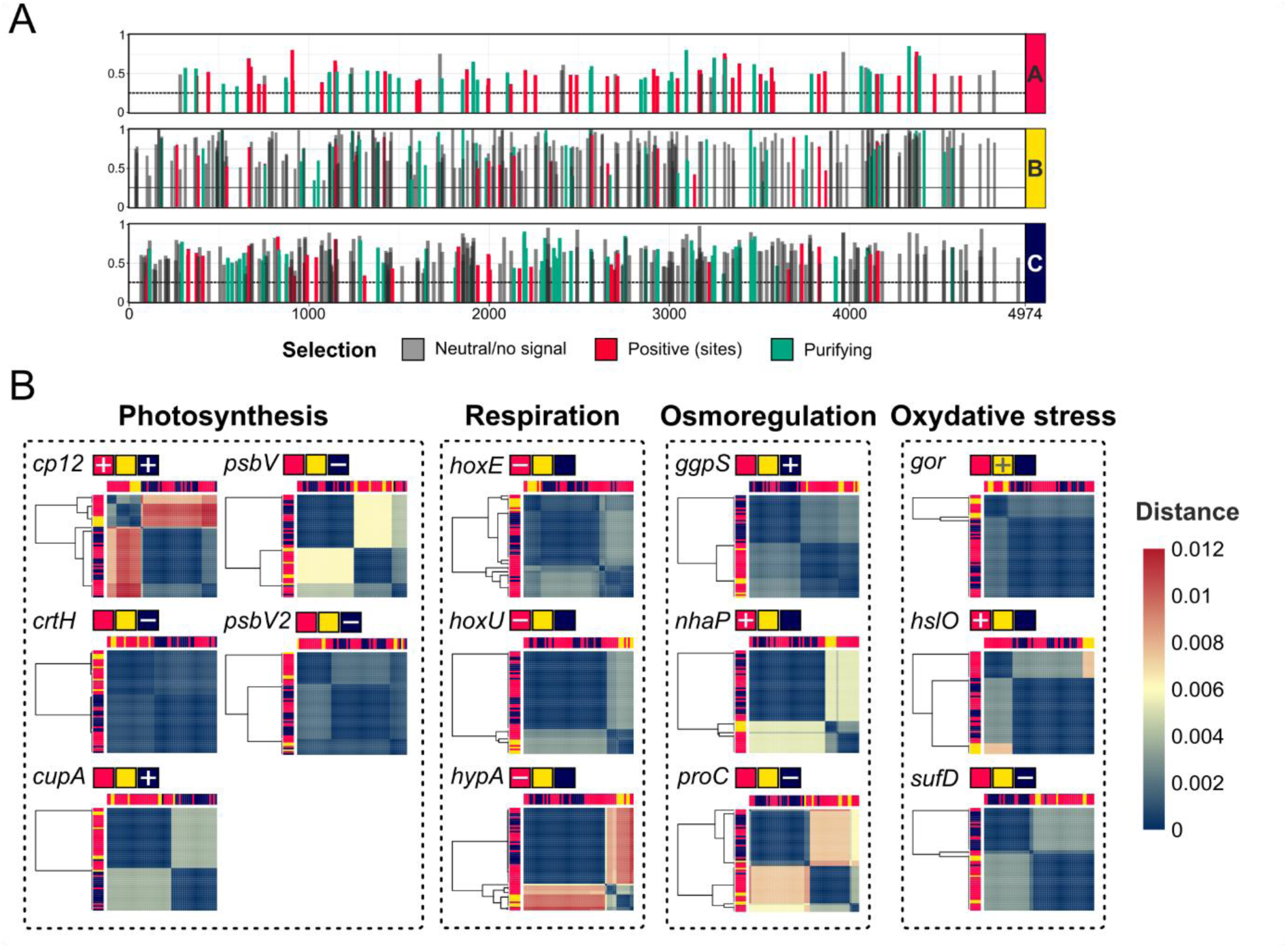
Genome-wide distribution and population structure of adaptively differentiated genes. **A** – Genome-wide distribution of adaptively differentiated genes. Bars indicate the positions of genes with significant differentiation throughout the composite reference genome, with bar height proportional to F_ST_ and colors indicating the inferred mode of selection. **B** – Heatmaps represent pairwise genomic distances among strains for adaptive genes. Colored bars indicate subpopulation assignments, with the inferred mode of selection associated with each subpopulation shown in squares (-: purifying selection, +: positive selection).

## Discussion

### Lake Dziani Dzaha as a model for reverse ecology

The microevolutionary processes shaping the population dynamics of continental bloom-forming cyanobacteria remain largely unresolved, in part because the very concept of microdiversity is confounded by taxonomic inconsistencies and the inherent difficulty of characterising niches in small shallow lake environments. Here, we leveraged the atypical features and history of Lake Dziani Dzaha, harboring a persistent and coherent *Limnospira platensis* population to investigate the spatial and temporal dimensions of microevolution. Although the niches colonized by *L. platensis* could not be comprehensively characterized *in situ*, the existence of objectively distinct macro-scale habitats allowed niche partitioning to be investigated in a reverse ecology perspective.

In addition to seasonal variations driven by precipitations and evaporation, which result in strong water-column stratification (Leboulanger *et al*. 2017, Sarazin *et al*. 2019), lake Dziani Dzaha has experience a major disruption of its water column since the onset of the seismic crisis. This one was characterized by a dramatic increase of CO_2_ pressure leading to a decline in pH and dissolved O_2_ concentrations, as well as enhanced dissolution of minerals from the surrounding magmatic rocks (Cadeau *et al*. 2022). Although the other macrohabitats (biofilms, stromatolites) are poorly characterized, they are expected to exhibit radically different physicochemical conditions. It is thought that the biotic green layers of stromatolites display high light specificity, as they harbour several cyanobacteria known for their adaptation to far-red light (Antonaru *et al*., 2025), and possess high-pH micro-niches near mineralizing cyanobacteria that promote carbonate precipitation (Merz, 1992, Riding, 2006, Jansson and Northen, 2010). Surface biofilms, in contrast, are known to exhibit pronounced microscale gradients in pH and concentration of oxygen (Walters *et al*., 2003, Wessel *et al*. 2014) and numerous solutes (*e.g.*, ammonium, hydrogen sulfide and carbon dioxide (Stewart and Franklin, 2008). Beyond abiotic contrasts, the three habitats would also harbor distinct microbial communities, as suggested by the divergent assemblages already reported between the water column and stromatolites (Gérard 2018).

### A highly coherent *Limnospira platensis* population displaying variable density through time

Despite the decline in population biomass recorded since monitoring began in 2010, and the dramatic decline triggered by the seismic event (May 2018 to November 2019), the *L. platensis* population of Lake Dziani Dzaha shows a remarkable degree of genetic cohesion across all habitats and sampling campaigns. All strains pairs shared an ANI value exceeding 99%, a threshold far above the classical used for species delineation (Richter & Rosseló-Móra, 2009) and which has been recently showed to discriminate *Microcystis* strains into highly metabolically coherent groups within/inside the species level. The genetic stability of the *Limnospira* population in Lake Dziani Dzaha is further supported by phylogenomic analyses, which show that strains isolated in 2024 consistently form robust monophyletic clusters with strains isolated in 2022 and 2010/2014. This pattern indicates that recent isolates are closely related to earlier members of the population and highlights the long-term coexistence of several temporally persistent lineages. The evolutionary characteristics of this population challenge the robust estimation of key recombination and demographic parameters (*e.g.*, μ, Nₑ, and δ) using dedicated tools (Didelot *et al*., 2007). In the absence of reliable estimates for these parameters, a parsimonious explanation for the persistence of most of genetically diverse lineages over the 14-year observation period is that mildly deleterious nonsynonymous substitutions are purged relatively slowly in this population, consistent with the nearly neutral theory of molecular evolution (Ohta *et al*., 1992).

### Gene loss/acquisition as dominant signatures of temporal adaptation

The transitional disequilibrium observed in 2022 for branch substitution rate and the recombination-initiation-to-mutation ratio is concomitant with a marked inflation of genome size and a reduction in coding sequence density. The doubling of insertion sequence (IS) copy number in 2022 genomes suggests that gene loss is at least partly attributable to IS-mediated recombination, which can cause complete gene deletion or partial inactivation, consistent with the concurrent accumulation of short pseudogenes (Touchon & Rocha, 2007). Such IS-driven genome rearrangements are consistent with a relaxation of the purifying selection that normally constrains IS proliferation, a phenomenon previously documented in bacteria colonizing new niches or undergoing population bottlenecks (Shapiro 1999a; Chain *et al*. 2004), and potentially favored here by the environmental stress imposed by the seismic event consequences and/or a drop in the effective *Limnospira* population size (Simonsen, 2022, Lynch, 2006, Bobay & Ochman, 2018).

Although gene loss has been associated with an increase in fitness (Koskiniemi et al., 2012; Albalat & Cañestro, 2016), this observation alone cannot allow to conclude to such associated adaptive benefits. However, the 2022 genomes show a specific enrichment in pair of genes encoding proteins of the chaperone-usher (CUP) system, involved in the assembly of fimbriae that play a role in cell adhesion and biofilm formation, and are frequently associated with surface colonization and persistence in harsh environments (Waksman & Hultgren, 2009; Busch & Waksman, 2012). These results, which are supported by the presence of *L. platensis* rafts structure in 2022 (see Fig. S1E), could therefore suggest that adaptation at this time was limited to protecting individuals from exposure to harmful physico-chemical conditions through an aggregated lifestyle, the accessory metabolic genes that would no longer confer significant fitness gains in such an environment being lost.

The genomic state of the 2022 population appears transitional rather than representative of a new evolutionary equilibrium. By 2024, gene content had already returned to a composition closely resembling that observed in pre-disturbance years (2010-2014), with 45 genes involved in signal transduction, macromolecular complexes regulation or cell envelope modification and polysaccharide biosynthesis, recovered within this two-year interval. Crucially, the widespread distribution of 2024 lineages across the phylogenetic tree (not exclusively in the vicinity of pre-2022 lineages), together with the diversity of the metabolic pathways in which they are involved, strongly suggest that these recoveries result from multiple independent soft gene sweep events, whereby beneficial gene variants spread rapidly through the population under positive selection, and considered as a major process in population evolutionary dynamics (Shapiro et al., 2012). Taken as a whole, these observations highlight a role for rare genes that goes beyond their implication in biotic interactions (Cordero & Polz, 2014). Although present in few lineages, these genes may provide a standing pool of functional diversity that can be rapidly recruited through soft sweeps, contributing to population resilience on remarkably short timescales.

### Niche partitioning: evidence from allelic variation

However, the gene presence/absence signal does not resolve spatial structure of the Limnospira population. No significant differentiation was detected between habitats or depths based on this metric, suggesting that large-scale gene content is primarily shaped by temporal rather than spatial selective pressures. The picture changes when allelic variation is considered, with three clusters of 2024 strains emerging as clearly distinct. F_ST_ patterns and the phylogenomic topology suggest that nsMC B and nsMC C independently diversified from nsMC A, with their differentiation maintained by contrasting selective pressures across the three populations. These selective pressures appear to be partly associated with differences in CO₂ acquisition, photosynthetic and respiratory metabolism, osmoregulation, and oxidative-stress responses, consistent with niche partitioning along functional traits expected to differ between the water column, stromatolites, and biofilms.

Since no cluster consists exclusively of strains isolated from a single habitat type, this hypothesis will require further validation, for instance by quantifying the relative abundance of each population across the three habitats. As observed for picocyanobacterial ecotypes, populations may be broadly distributed across habitats while differing markedly in their relative abundances (Biller *et al*., 2014). In Lake Dziani Dzaha, exchange between habitats may also contribute to this pattern, particularly because acidification and the resulting decrease in pH can damage stromatolite structures and promote the release and redistribution of associated populations.

### The complexity of microevolutionary modes

Two non-exclusive models have been proposed to explain the microdiversity. The Renaissance model posits that horizontal gene transfer (HGT) is the primary driver of ecological differentiation, with novel trait acquisitions enabling access to new resources and qualitatively distinct niches (Larkin & Martiny, 2017). By contrast, the Maestro model emphasizes that point mutations accumulating along phylogenetic lineages as the foundation of adaptive divergence to sympatric partition based on functional trait optimization (Larkin & Martiny, 2017). Our results confirm that both models are operating simultaneously within *L. platensis* population of lake Dziani Dzaha. Interestingly, new habitats colonization by strains originating from the water column is more consistent with a “Maestro”-type divergence, which has been linked to the differentiation among ecotypes following abiotic parameters such as temperature, pH or size of associated-particles (Doré *et al*., 2020, Newton *et al*., 2007, Shapiro *et al*., 2012). If the niches underlying this differentiation were associated with stromatolites and biofilms, this would suggest that a fine-tuning of central metabolic pathways is sufficient for *Limnospira* to adapt to these supposedly contrasting environments. However, as illustrated by these previous examples, the ecological differentiation underlying these genomic patterns may also occur within the water column itself, among microhabitats that remain to be characterized. In contrast, a ‘Renaissance’-type dynamics appear in this system to be associated with population resilience after a strong environmental perturbation rather than with stable niche colonization. Evolution experiments have shown the bacterial capacity to adapt to rapid shifts to high temperature or high CO_2_ concentration, within a range of few months or 2-4 years, respectively (Kent *et al*., 2018, Hutchins *et al*., 2015). But, to our knowledge, our study is the first to document, in a natural system, such rapid genome-level responses to environmental change: an initial phase of genome streamlining and a shift in lifestyle, followed by genome expansion and the near-complete recovery of adaptive genes.

## Acknowledgements

Authors thank the Air Austral Airline Company and “Le Relais Dziani” in Mayotte for their assistance and support. The field permit was granted by the Conservatoire du Littoral et des Rivages Lacustres, Antenne Océan Indien.

---

**Figure S1.**
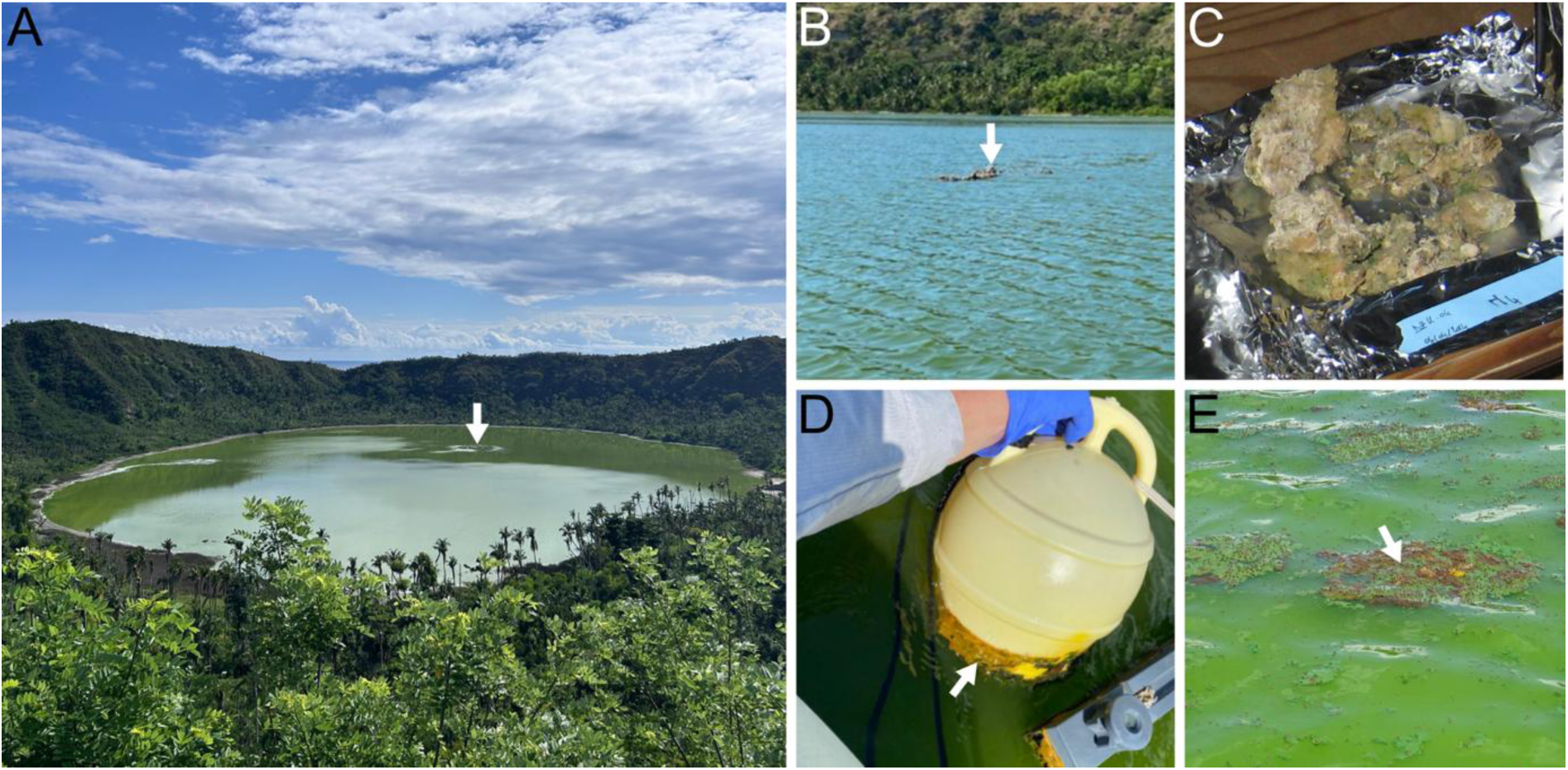
Sampling field overview, stromatolites, biofilm and cyanobacterial aggregates in Lake Dziani Dzaha. **A** - DZ25-11 panoramic view of Lake Dziani Dzaha from the east crest of the crater. ©A. Le Vern, MNHN. White arrow showing CO_2_ bubbles rising from the pit. **B –** In situ photography of stromatolite S3 emerging from the lakes surface (white arrow) during the DZ24-06 sampling campaign. ©C. Bernard, MNHN. **C** - Photography of stromatolite S4 back at the field lab from the DZ14-04 sampling campaign. ©C. Bernard, MNHN. **D** - DZ24-06 encrusted biofilm (white arrow) on the surface of a buoy (B1 sampling site). **E –** Floating cyanobacterial aggregates - ‘rafts’ (white arrow) - observed in 2022, characterized by the absence of extracellular matrix, cyanobacteria easily dissociating from others upon disturbance. ©D. Jézéquel, IPGP.

**Figure S2.**
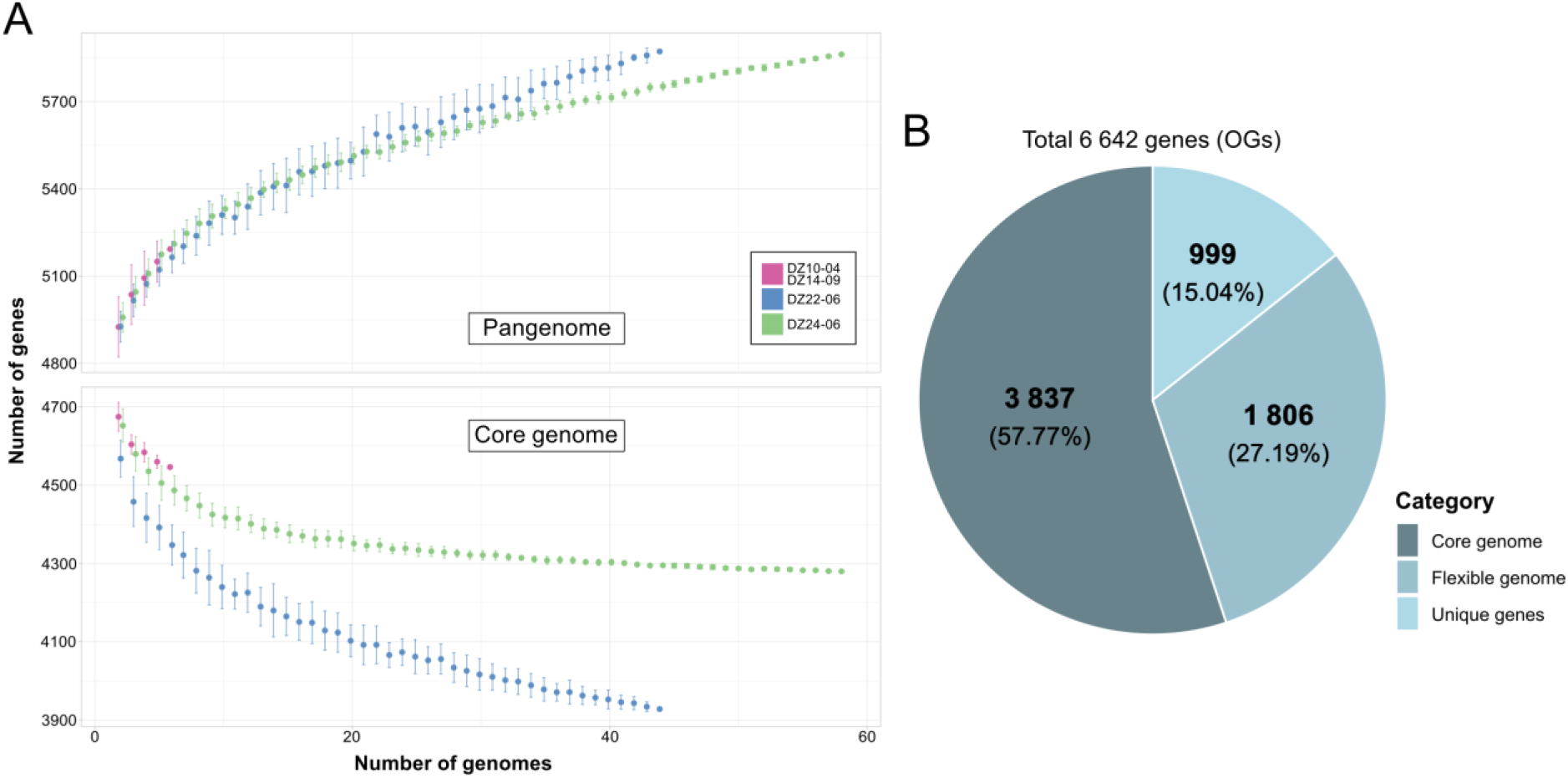
Pangenome structure : saturation and proportions. **A** - Rarefaction curves for the pangenome (top) and the core genome (bottom) as a function of the number of genomes, by year of strain collection: 2010 and 2014 (pink), 2022 (blue), and 2024 (green). **B** - Distribution of the 6,642 orthologous groups (OGs) identified in the pangenome, divided into three categories: core genome (dark gray; 3,837 OGs, 57.77%), unique genes (light blue; 999 OGs, 15.04%), and the rest of the flexible genome (medium blue-gray; 1,806 OGs, 27.19%).

**Figure S3.**
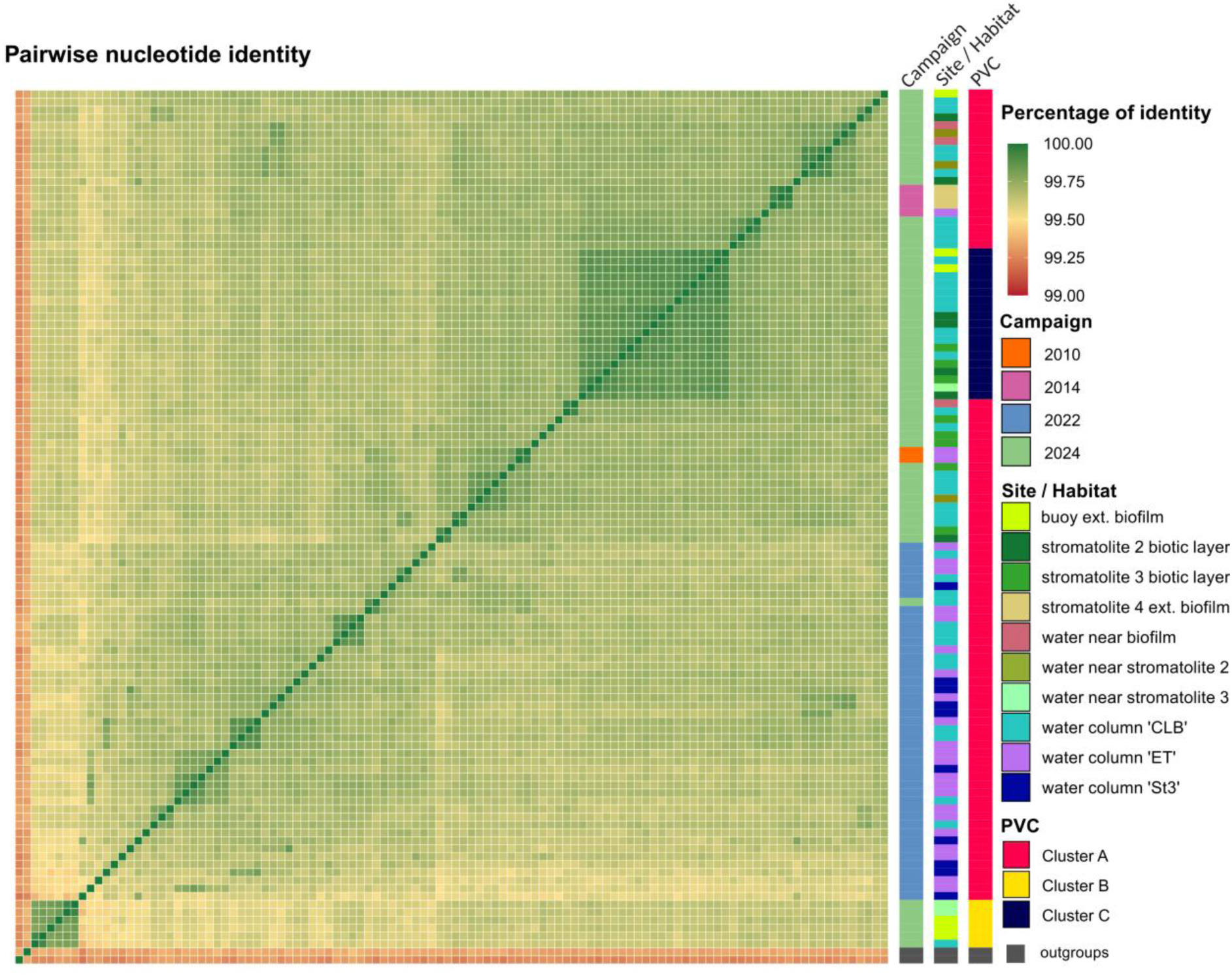
Pairwise average nucleotide identity (ANI) between all studied strains. Heatmap representing the percentage of identity between shared sequences of all studied strains and outgroup strains (SAJ85.79 and TJSD092). Annotations bars are colored as a function of the sampling campaign the strain has been isolated from, the site/habitat where the strain were sampled, and the nsmC (nonsynonymous mutation cluster).

**Figure S4.**
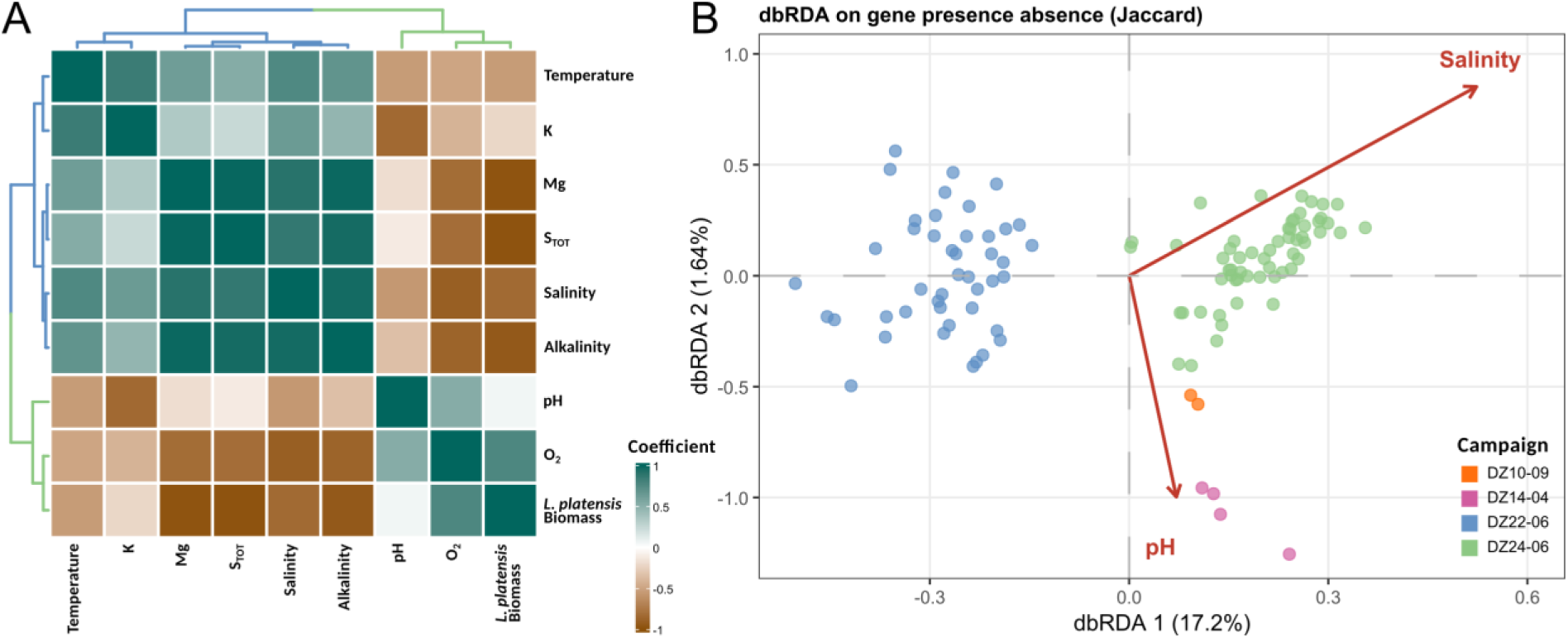
Environmental drivers of Limnospira platensis flexible genome composition. A - Spearman’s collinearity heatmap representing the links between environmental parameters. Dendrograms reflect the hierarchical clustering of the colinear variable. B – distance-based redundancy analysis (db-RDA) performed on a Jaccard dissimilarity matrix across all sampling campaigns. Salinity and pH act as explanatory proxies of the variables they are colinear with. Vector arrows represent the weight and directional influence that the environmental variables have on the flexible genome structure.

## References

Aiba S, Ogawa T. Assessment of Growth Yield of a Blue—green Alga, Spirulina platensis, in Axenic and Continuous Culture. Microbiology 1977;102:179–182. 10.1099/00221287-102-1-179

Albalat R, Cañestro C. Evolution by gene loss. Nat Rev Genet 2016;17:379–391. 10.1038/nrg.2016.39

Antonaru LA et al. Evolution of far-red light photoacclimation in cyanobacteria. Current Biology 2025;35:2539–2553.e4. 10.1016/j.cub.2025.04.038

Banerji A et al. Evaluating putative ecological drivers of microcystin spatiotemporal dynamics using metabarcoding and environmental data. Harmful Algae 2019;86:84–95. 10.1016/j.hal.2019.05.004

Bankevich A et al. SPAdes: a new genome assembly algorithm and its applications to single-cell sequencing. J Comput Biol 2012;19:455–477. 10.1089/cmb.2012.0021

Bernard C et al. Very Low Phytoplankton Diversity in a Tropical Saline-Alkaline Lake, with Co-dominance of Arthrospira fusiformis (Cyanobacteria) and Picocystis salinarum (Chlorophyta). Microb Ecol 2019;78:603–617. 10.1007/s00248-019-01332-8

Biller SJ et al. Genomes of diverse isolates of the marine cyanobacterium Prochlorococcus. Sci Data 2014;1:140034. 10.1038/sdata.2014.34

Biller SJ et al. Prochlorococcus: the structure and function of collective diversity. Nat Rev Microbiol 2015;13:13–27. 10.1038/nrmicro3378

Bobay L-M. The Prokaryotic Species Concept and Challenges. In: Tettelin H, Medini D (eds), The Pangenome: Diversity, Dynamics and Evolution of Genomes. Cham (CH): Springer, 2020.

Bobay L-M, Ochman H. Factors driving effective population size and pan-genome evolution in bacteria. BMC Evol Biol 2018;18:153. 10.1186/s12862-018-1272-4

Bolger AM, Lohse M, Usadel B. Trimmomatic: a flexible trimmer for Illumina sequence data. Bioinformatics 2014;30:2114–2120. 10.1093/bioinformatics/btu170

Briand E et al. Spatiotemporal changes in the genetic diversity of a bloom-forming *Microcystis aeruginosa* (cyanobacteria) population. The ISME Journal 2009;3:419–429. 10.1038/ismej.2008.121

Busch A, Waksman G. Chaperone–usher pathways: diversity and pilus assembly mechanism. Philos Trans R Soc Lond B Biol Sci 2012;367:1112–1122. 10.1098/rstb.2011.0206

Cadeau P et al. Impact of the seismo-volcanic crisis offshore Mayotte on the Dziani Dzaha Lake. Comptes Rendus Géoscience 2022;354:299–316. 10.5802/crgeos.172

Cadeau P et al. Carbon isotope evidence for large methane emissions to the Proterozoic atmosphere. Sci Rep 2020;10:18186. 10.1038/s41598-020-75100-x

Chain PSG et al. Insights into the evolution of *Yersinia pestis* through whole-genome comparison with *Yersinia pseudotuberculosis*. Proc Natl Acad Sci USA 2004;101:13826–13831. 10.1073/pnas.0404012101

Charrad M et al. NbClust: An R Package for Determining the Relevant Number of Clusters in a Data Set. Journal of Statistical Software 2014;61:1–36. 10.18637/jss.v061.i06

Chklovski A et al. CheckM2: a rapid, scalable and accurate tool for assessing microbial genome quality using machine learning. Nat Methods 2023;20:1203–1212. 10.1038/s41592-023-01940-w

Cordero OX, Polz MF. Explaining microbial genomic diversity in light of evolutionary ecology. Nat Rev Microbiol 2014;12:263–273. 10.1038/nrmicro3218

Croucher NJ et al. Rapid phylogenetic analysis of large samples of recombinant bacterial whole genome sequences using Gubbins. Nucleic Acids Research 2015;43:e15–e15. 10.1093/nar/gku1196

De Cáceres M, Legendre P. Associations between species and groups of sites: indices and statistical inference. Ecology 2009;90:3566–3574. 10.1890/08-1823.1

Didelot X, Falush D. Inference of bacterial microevolution using multilocus sequence data. Genetics 2007;175:1251–1266. 10.1534/genetics.106.063305

Didelot X, Wilson DJ. ClonalFrameML: efficient inference of recombination in whole bacterial genomes. PLoS Comput Biol 2015;11:e1004041. 10.1371/journal.pcbi.1004041

Doré H et al. Evolutionary Mechanisms of Long-Term Genome Diversification Associated With Niche Partitioning in Marine Picocyanobacteria. Front Microbiol 2020;11. 10.3389/fmicb.2020.567431

Emms DM, Kelly S. OrthoFinder: phylogenetic orthology inference for comparative genomics. Genome Biol 2019;20:238. 10.1186/s13059-019-1832-y

Feuillet N et al. Birth of a large volcanic edifice offshore Mayotte via lithosphere-scale dyke intrusion. Nature Geoscience 2021;14:1–9. 10.1038/s41561-021-00809-x

Flombaum P et al. Present and future global distributions of the marine Cyanobacteria Prochlorococcus and Synechococcus. Proceedings of the National Academy of Sciences of the United States of America 2013;110. 10.1073/pnas.1307701110

Fuhrman JA, Campbell L. Microbial microdiversity. Nature 1998;393:410–411.

Gérard E et al. Key Role of Alphaproteobacteria and Cyanobacteria in the Formation of Stromatolites of Lake Dziani Dzaha (Mayotte, Western Indian Ocean). Front Microbiol 2018;9:796. 10.3389/fmicb.2018.00796

Grébert T et al. Light color acclimation is a key process in the global ocean distribution of Synechococcus cyanobacteria. Proceedings of the National Academy of Sciences 2018;115:E2010–E2019. 10.1073/pnas.1717069115

Halary S et al. Metagenome-Based Exploration of Bacterial Communities Associated with Cyanobacteria Strains Isolated from Thermal Muds. Microorganisms 2022;10:2337. 10.3390/microorganisms10122337

Halary S et al. Intra-population genomic diversity of the bloom-forming cyanobacterium, Aphanizomenon gracile, at low spatial scale. ISME Commun 2023;3:57. 10.1038/s43705-023-00263-3

Hamlaoui S et al. The Culture Collection of Cyanobacteria and Microalgae at the French National Museum of Natural History: A Century Old But Still Alive and Kicking! Including in Memoriam: Professor Alain Couté. crya 2022;43:41–83. 10.5252/cryptogamie-algologie2022v43a3

Huerta-Cepas J et al. eggNOG 5.0: a hierarchical, functionally and phylogenetically annotated orthology resource based on 5090 organisms and 2502 viruses. Nucleic Acids Research 2019;47:D309–D314. 10.1093/nar/gky1085

Huré A et al. Metabolite diversity of Microcystis strains shows tight correspondence to genotype and may contribute to ecotype specificities. Commun Biol 2026;9:305. 10.1038/s42003-026-09599-7

Hutchins DA et al. Irreversibly increased nitrogen fixation in Trichodesmium experimentally adapted to elevated carbon dioxide. Nat Commun 2015;6:8155. 10.1038/ncomms9155

Ibelings B, Mur L, Walsby A. Diurnal changes in buoyancy and vertical distribution in populations of Microcystis in two shallow lakes. Journal of Plankton Research 1991;13:419–436.

Jackrel SL et al. Genome evolution and host-microbiome shifts correspond with intraspecific niche divergence within harmful algal bloom-forming Microcystis aeruginosa. Mol Ecol 2019;28:3994–4011. 10.1111/mec.15198

Jain C et al. High throughput ANI analysis of 90K prokaryotic genomes reveals clear species boundaries. Nat Commun 2018;9:5114. 10.1038/s41467-018-07641-9

Johnson ZI et al. Niche Partitioning Among Prochlorococcus Ecotypes Along Ocean-Scale Environmental Gradients. Science 2006;311:1737–1740. 10.1126/science.1118052

Katoh K, Standley DM. MAFFT Multiple Sequence Alignment Software Version 7: Improvements in Performance and Usability. Mol Biol Evol 2013;30:772–780. 10.1093/molbev/mst010

Kent AG, Garcia CA, Martiny AC. Increased biofilm formation due to high temperature adaptation in marine Roseobacter. Nat Microbiol 2018;3:989–995. 10.1038/s41564-018-0213-8

Koskiniemi S et al. Selection-driven gene loss in bacteria. PLoS Genet 2012;8:e1002787. 10.1371/journal.pgen.1002787

Kuijpers MCM et al. Intraspecific divergence within *Microcystis aeruginosa* mediates the dynamics of freshwater harmful algal blooms under climate warming scenarios. Proc R Soc B 2025;292:20242520. 10.1098/rspb.2024.2520

Larkin AA, Martiny AC. Microdiversity shapes the traits, niche space, and biogeography of microbial taxa. Environmental Microbiology Reports 2017;9:55–70. 10.1111/1758-2229.12523

Leboulanger C et al. Microbial Diversity and Cyanobacterial Production in Dziani Dzaha Crater Lake, a Unique Tropical Thalassohaline Environment. PLOS ONE 2017;12:e0168879. 10.1371/journal.pone.0168879

Lesturgie P, Planes S, Mona S. Coalescence times, life history traits and conservation concerns: An example from four coastal shark species from the Indo-Pacific. 2022.

Lynch M. Streamlining and simplification of microbial genome architecture. Annu Rev Microbiol 2006;60:327–349. 10.1146/annurev.micro.60.080805.142300

Martiny AC et al. Taxonomic resolution, ecotypes and the biogeography of Prochlorococcus. Environmental Microbiology 2009;11:823–832. 10.1111/j.1462-2920.2008.01803.x

Mattei F et al. Chromatic acclimation shapes phytoplankton biogeography. Science Advances 2025;11:eadr9609. 10.1126/sciadv.adr9609

Menden-Deuer S, Lessard EJ. Carbon to volume relationships for dinoflagellates, diatoms, and other protist plankton. Limnology and Oceanography 2000;45:569–579. 10.4319/lo.2000.45.3.0569

Merz MUE. Biology of carbonate precipitation by cyanobacteria. Facies 1992;29:80–80. 10.1007/BF02536920

Minh BQ et al. IQ-TREE 2: New Models and Efficient Methods for Phylogenetic Inference in the Genomic Era. Mol Biol Evol 2020;37:1530–1534. 10.1093/molbev/msaa015

Moore L, Goericke R, Chisholm S. Comparative physiology of Synechococcus and Prochlorococcus: Influence of light and temperature on growth, pigments, fluorescence and absorptive properties. Marine Ecology-progress Series - MAR ECOL-PROGR SER 1995;116:259–275. 10.3354/meps116259

Newton RJ et al. Phylogenetic Ecology of the Freshwater Actinobacteria acI Lineage. Applied and Environmental Microbiology 2007;73:7169–7176. 10.1128/AEM.00794-07

O’Neil JM et al. The rise of harmful cyanobacteria blooms: The potential roles of eutrophication and climate change. Harmful Algae 2012;14:313–334. 10.1016/j.hal.2011.10.027

Ohta T. The Nearly Neutral Theory of Molecular Evolution. Annual Review of Ecology and Systematics 1992;23:263–286.

Oksanen J et al. vegan: Community Ecology Package. 2026. 2026.

R Core Team. R: A Language and Environment for Statistical Computing.

Richter M, Rosselló-Móra R. Shifting the genomic gold standard for the prokaryotic species definition. Proc Natl Acad Sci U S A 2009;106:19126–19131. 10.1073/pnas.0906412106

Riding R. Cyanobacterial calcification, carbon dioxide concentrating mechanisms, and Proterozoic– Cambrian changes in atmospheric composition. Geobiology 2006;4:299–316. 10.1111/j.1472-4669.2006.00087.x

Rocap G et al. Genome divergence in two Prochlorococcus ecotypes reflects oceanic niche differentiation. Nature 2003;424:1042–1047. 10.1038/nature01947

Roussel T et al. Monospecific renaming within the cyanobacterial genus Limnospira (Spirulina) and consequences for food authorization. J Appl Microbiol 2023;134:lxad159. 10.1093/jambio/lxad159

Sarazin G et al. Geochemistry of an endorheic thalassohaline ecosystem: the Dziani Dzaha crater lake (Mayotte Archipelago, Indian Ocean). Comptes Rendus Géoscience 2021;352:559–577. 10.5802/crgeos.43

Schuergers N, Wilde A. Appendages of the Cyanobacterial Cell. Life (Basel) 2015;5:700–715. 10.3390/life5010700

Seemann T. Prokka: rapid prokaryotic genome annotation. Bioinformatics 2014;30:2068–2069. 10.1093/bioinformatics/btu153

Seemann T. tseemann/snippy. 2015. 2015.

Shapiro BJ et al. Population Genomics of Early Events in the Ecological Differentiation of Bacteria. Science 2012;336:48–51. 10.1126/science.1218198

Shapiro JA. Genome system architecture and natural genetic engineering in evolution. Ann N Y Acad Sci 1999;870:23–35. 10.1111/j.1749-6632.1999.tb08862.x

Simonsen AK. Environmental stress leads to genome streamlining in a widely distributed species of soil bacteria. ISME J 2022;16:423–434. 10.1038/s41396-021-01082-x

Stewart PS, Franklin MJ. Physiological heterogeneity in biofilms. Nat Rev Microbiol 2008;6:199–210. 10.1038/nrmicro1838

Suyama M, Torrents D, Bork P. PAL2NAL: robust conversion of protein sequence alignments into the corresponding codon alignments. Nucleic Acids Res 2006;34:W609–W612. 10.1093/nar/gkl315

Syberg-Olsen MJ et al. Pseudofinder: Detection of Pseudogenes in Prokaryotic Genomes. Mol Biol Evol 2022;39:msac153. 10.1093/molbev/msac153

Touchon M, Rocha EPC. Causes of insertion sequences abundance in prokaryotic genomes. Mol Biol Evol 2007;24:969–981. 10.1093/molbev/msm014

Waksman G, Hultgren SJ. Structural biology of the chaperone–usher pathway of pilus biogenesis. Nat Rev Microbiol 2009;7:765–774. 10.1038/nrmicro2220

Walters MC et al. Contributions of antibiotic penetration, oxygen limitation, and low metabolic activity to tolerance of Pseudomonas aeruginosa biofilms to ciprofloxacin and tobramycin. Antimicrob Agents Chemother 2003;47:317–323. 10.1128/AAC.47.1.317-323.2003

Wessel AK et al. Oxygen limitation within a bacterial aggregate. mBio 2014;5:e00992. 10.1128/mBio.00992-14

Wu H et al. Selection of photosynthetic traits by turbulent mixing governs formation of cyanobacterial blooms in shallow eutrophic lakes. The ISME Journal 2024;18:wrae021. 10.1093/ismejo/wrae021

Wu Y et al. Seasonal dynamics of water bloom-forming *Microcystis* morphospecies and the associated extracellular microcystin concentrations in large, shallow, eutrophic Dianchi Lake. Journal of Environmental Sciences 2014;26:1921–1929. 10.1016/j.jes.2014.06.031

Xie Z, Tang H. ISEScan: automated identification of insertion sequence elements in prokaryotic genomes. Bioinformatics 2017;33:3340–3347. 10.1093/bioinformatics/btx433

Yéprémian C et al. Microcystin ecotypes in a perennial *Planktothrix agardhii* bloom. Water Research 2007;41:4446–4456. 10.1016/j.watres.2007.06.028

Zhaxybayeva O et al. Intertwined Evolutionary Histories of Marine Synechococcus and Prochlorococcus marinus. Genome Biol Evol 2009;1:325–339. 10.1093/gbe/evp032

Zinke J, Reijmer JJG, Thomassin BA. Systems tracts sedimentology in the lagoon of Mayotte associated with the Holocene transgression. Sedimentary Geology 2003;160:57–79. 10.1016/S0037-0738(02)00336-6

Collection de cyanobactéries et de microalgues vivantes (PMC-ALCP) | Molécules de Communication et Adaptation des Micro-organismes (MCAM). https://mcam.mnhn.fr/fr/collection-de-cyanobacteries-et-de-microalgues-vivantes-pmc-alcp-470.

